# The *foraging* gene affects alcohol sensitivity, metabolism and memory in *Drosophila*

**DOI:** 10.1101/2021.02.09.430533

**Authors:** Anne S. Oepen, Jamie Catalano, Reza Azanchi, Karla R. Kaun

## Abstract

The genetic basis of alcohol use disorder (AUD) is complex. Understanding how natural genetic variation contributes to alcohol phenotypes can help identify mechanisms underlying the genetic contribution of AUD. Recently, a single nucleotide polymorphism in the human foraging (*for*) gene ortholog, Protein Kinase cGMP-Dependent 1 (PRKG1), was found to be associated with stress-induced risk for alcohol abuse. However, the mechanistic role that PRKG1 plays in AUD is not well understood. We use natural variation in the *Drosophila for* gene to describe how variation of cGMP-dependent protein kinase (PKG) activity modifies ethanol-induced phenotypes. We found that variation in *for* affects ethanol-induced increases in locomotion and memory of the appetitive properties of ethanol intoxication. Further, these differences may stem from the ability to metabolize ethanol. Together, this data suggests that natural variation in PKG modulates cue reactivity for alcohol, and thus could influence alcohol cravings by differentially modulating metabolic and behavioral sensitivities to alcohol.

## Introduction

Determining the mechanisms through which individual genes influence natural variation in behavior has been difficult due to the complexity of the genetic basis of heritable behavior. A notable exception to the genetic complexity underlying behavior is the foraging (*for*) gene in *Drosophila* melanogaster (1). Variants of *for* show increased (rovers or *for^R^*) or decreased (sitters or *for^s^*) cGMP-dependent protein kinase (PKG) activity, which affects food search behavior (2). Rovers show increased pathlength while foraging, whereas sitters show decreased pathlength and stay longer at one food source (1, 3) *for* also causes phenotypical pleiotropy in flies, in part due to multiple promotors driving expression in different cell types (4), and at different developmental stages (5–7). The foraging gene affects foraging behavior (1, 8–13)), fat and glucose stores (4, 14), food intake (14, 15), sucrose response (16, 17), sleep (18), habituation (17, 19, 20), nociception (21), oviposition (22), stress response (23–25), as well as learning and memory (17, 20, 26–30)

Moreover, for’s role as a behavioral modifier in a wide range of behaviors is conserved across species including Apis mellifera (31), Xenopus laevis (32), Mus musculus (33), Caenorhabditis elegans (34, 35), and Homo sapiens (36, 37). Given for’s conserved role, understanding the mechanistic actions through which *for* affects behavior may lead to insight regarding conserved functions of how natural genetic variation alters behavior.

More recently, Protein Kinase cGMP-Dependent 1 (PRKG1), the human ortholog of *for* (36, 38), was implicated in stress-induced risk for alcohol abuse (39, 40). A gene-by-environment genome-wide interaction study (GEWIS) investigating mechanisms by which traumatic life events influence genetic variation in relation to alcohol misuse, revealed several risk alleles for alcohol use disorder (AUD) including PRKG1 (40). Similarly, a genome-wide association study (GWAS) implicated PRKG1 in other stressed induces phenotypes like post-traumatic stress disorder (41).

Other components of cGMP signaling have also been implicated in alcohol-associated phenotypes. Consumption of alcohol increases cGMP levels in the rat cortex, striatum and hippocampus (42). Increasing cGMP levels in the rat ventral tegmental area (VTA) or medial prefrontal cortex (mPFC) reduces the ability of alcohol deprivation to enhance drinking, which is reversed by inhibiting PKG (43). Finally, deletion of PKG type II in mice reduces alcohol’s sedative effects and increases alcohol consumption (44). NO/cGC/cGMP/PKG signaling also causes neuroadaptive changes in synaptic activity, thereby affecting distinct forms of learning and memory, such as object recognition, motor adaptation and fear conditioning (45–48). This signaling pathway similarly inhibits dopamine release in brain regions that are involved in addiction (49–51), and contributes to cocaine self-administration (52) and morphine induced neuroadaptation (53). This compounding evidence suggests PRGK1 plays a critical role in alcohol consumption, and calls for a better understanding of *for*’s mechanistic role in alcohol induced behaviors.

*Drosophila* has proven to be a valuable model organism for identifying genes and elucidating mechanisms associated with AUD (54–56). In addition to genomic, transcriptomic and proteomic approaches, simple forward and reverse genetic approaches can be performed to identify genes that affect alcohol-induced behaviors, and to elucidate the cellular and molecular mechanisms underlying candidate genes linked to AUD. Due to the huge variety of genetic tools available in *Drosophila*, many cell types, neural circuits and genes have been linked to an alcohol phenotype (57–73). Therefore, investigating *for*’s role in relation to alcohol phenotypes in the fly might help to unravel the function of PRKG1 in alcohol addiction in humans.

## Material and Methods

### Fly Stocks and Rearing Conditions

The following fly strains were used: *for^s^* (sitter), *for^R^* (rover) and *for^s2^. for^s^* and *for^R^* flies carry the natural rover/sitter polymorphisms in for. *for^s2^* flies have a rover genetic background but carry a gamma radiation-induced mutation in *for* that results in lower PKG activity levels and sitter behavior in many of the *for*-related phenotypes (2, 10, 11). However, the *for^s2^* mutant does not always affect *for*-dependent phenotypes, and to our knowledge, specific alterations in DNA sequence as a result of the s2 mutation are unknown (74, 75). Flies were raised on standard cornmeal agar food media with tegosept anti-fungal agent. Flies were kept at 24°C and 65% humidity with a 14/10-hour light/dark cycle. Male flies for all strains were collected 1-2 days after hatching and used for behavioral experiments at day 3-5 after eclosion.

### Group Activity

In order to analyze the ethanol-induced changes in activity, we used a video based behavioral apparatus and software that enables an automatic quantification of group locomotion activity in *Drosophila* (70) (Fig 1A). The fly Group Activity Monitor (FlyGrAM) consists of four circular arenas that are all individually connected to an air and vacuum source to allow for a constant airflow (Fig 1A,B). Each arena is 37 mm diameter, 2.5 mm tall, and covered with a clear acrylic sheet allowing the flies to walk freely in the arena, but preventing them from flying. Ten male flies (*for^s^, for^R^* or *for^s2^*) were placed in each circular arena and placed at 24°C for 10 minutes to provide time for flies to adapt to the arena environment. Next, the arena with the flies was placed in the tracking apparatus, and flies were provided a constant humidified airflow of 115 units (1856 ml/min) for 15 minutes, which allowed their locomotion to decrease to a stable baseline (Fig 1A).

**Fig. 1.**
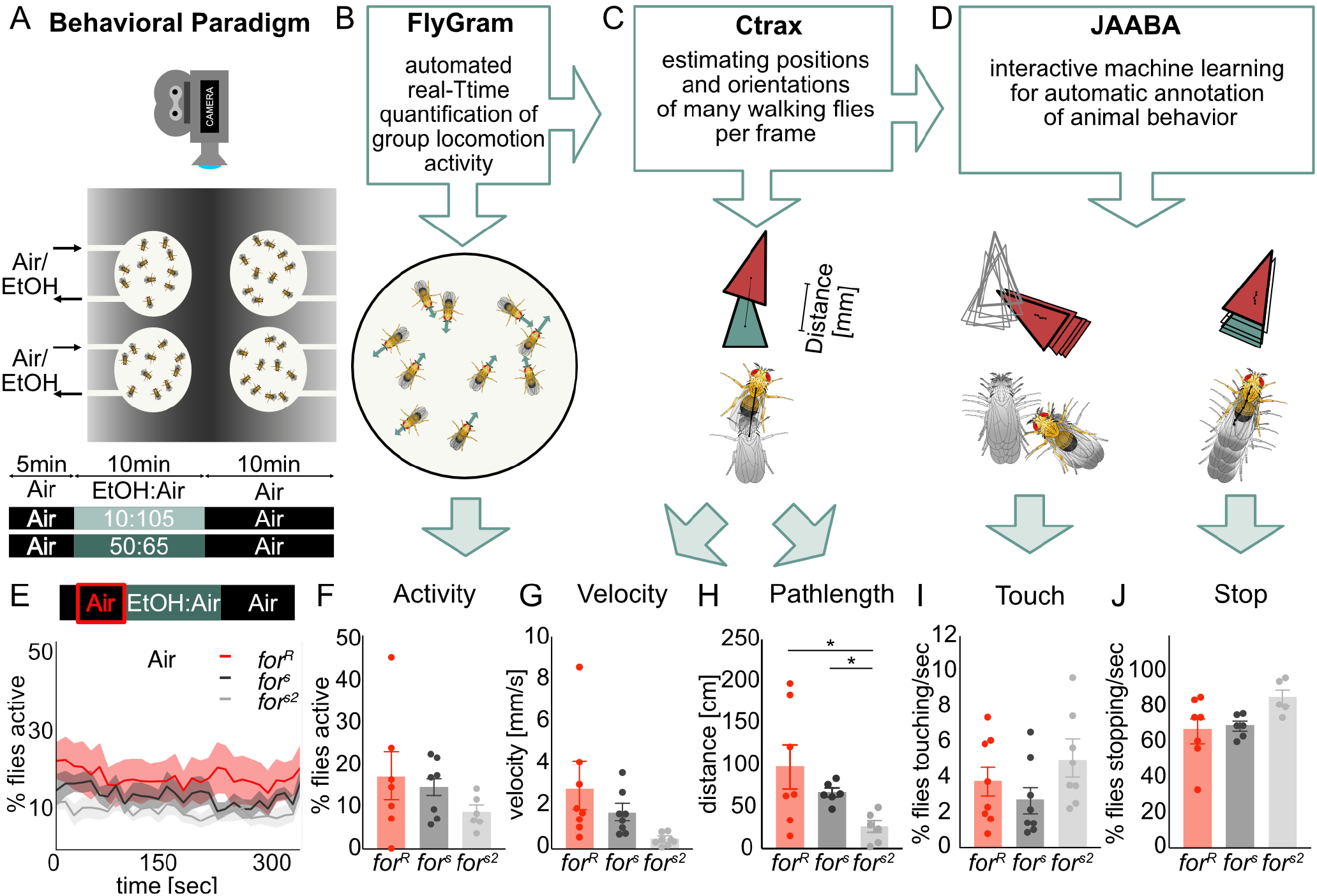
*for* does not affect spontaneous open field behavior. A) The flyGrAM arena consists of 4 circular arenas each filled with 10 flies of different strains. Flies were tracked while being given humidified air for 5min, ethanol for 10min, then humidified air for 5min. (B) Group activity within each arena was analyzed with the FlyGrAM software while recording the flies during the behavioral paradigm. (C) Ctrax was used to estimate the position and orientation of the 10 flies in each arena per frame. (D) The interactive machine learning for automatic annotation of animal behavior (JAABA) was fed per-frame feature information from ctrax to reveal information about social interaction of the flies. (E,F) Lines depict mean+/- standard error. (F-J) Graphs show mean+/- standard error. The spontaneous activity levels of flies given humidified air remain at 15% and show no significant difference between strains. (G) Velocity of flies given humidified air for 5min is between 1-3mm/s with no significant difference between strains. (H) Flies cover distances between 30-100cm in five minutes, with no significant differences between strains. (I) Less than 5% of the flies touch per second when given humidified air, with no significant differences between strains. (J) More than 60% of the flies show stopping behavior per second when given humidified air, with no significant differences between strains.

Ten flies were gently placed via mouth pipette into each arena, and initially provided humidified airflow for five minutes to gain a measurement of baseline activity. Subsequently, vaporized ethanol was introduced to the flies in the arena by changing the gas flow-through to a mix of ethanol:air ratio (high concentration: 50:65, low concentration: 10:105) for 10 minutes. Finally, the airflow was shifted back to 100% air for 10 minutes (Fig 1A). The group activity of the flies are automatically recorded for the entire duration (70). Each experiment was replicated 8 times (n=8 comprises 8 groups of 10 flies per group, thus 80 flies). In order to ensure no strain-specific effects were due to a single arena and no spatial bias of placement within the apparatus, the strains tested were counterbalanced across arenas. Experiments were run at a steady temperature around 24°C ± 1.5°C in a dark chamber to reduce the influence of visual cues on group activity.

### High Content Tracking

FlyGrAM videos were analyzed using Ctrax (Branson et al., 2009) to extract information about individual flies’ position, orientation and trajectories (Fig 1B). The output of Ctrax tracking is an ellipse fit to each fly within an arena throughout the experiment. It is defined by the centroid, the fly’s orientation as well as the length of two body axis, the minor and major body axis, needed for the calculation of the perframe features. For example, velocity is defined as the speed of the center of rotation, being the point on the fly that translates the least from one frame to the next. The velocity is calculated by the magnitude of the vector between the fly’s center of rotation in frames t 1 to t, normalized by the frame rate (76). This allowed us to gain information about the velocity of the flies per frame. Total distance travelled (pathlength) before, during and after ethanol exposures was calculated as the integral of speed, i.e. summing the perframe (30fps) velocities (mm/s) measured during 5 minutes. JAABA Classifiers (77) were trained on the Ctrax perframe features to gain information about specific social and locomotion behaviors: Touch and Stop (Fig 1C). The JAABA learning algorithm is fed with specific pre-defined behavioral classifiers in order to scan across all frames for the classified behavior. In this study we made use of the existing classifiers for stopping and touching behavior introduced by Branson and colleagues (77). To further analyze the effect of ethanol on these behavioral features, the data was split into 4 phases each lasting 5 minutes, baseline(0-5min), early ethanol(5-10min), late ethanol(10-15min) and recovery(15-20min). The Stop and Touch data was normalized to the baseline behavior to detect changes in these features caused by alcohol exposure.

### Memory for Ethanol Reward

To test whether *for* affected the ability to associate a cue with ethanol, we exposed *for^R^, for^s^* and *for^s2^* flies to two consecutive odor cues (1:36 isoamyl alcohol or ethyl acetate in mineral oil) with the second odor paired with an intoxicating dose of 60% ethanol vapor (90:60 ethanol:air) (78, 79). Flies were exposed to 10 min of the first odor followed by 10 min of the second odor, which is paired with 60% ethanol vapor. Flies were trained three times with 50 min breaks between each session. Flies were placed in perforated 14mL canonical tubes with mesh lids, and placed in 30cmx15cmx15cm training boxes with passive vacuum while being exposed to odors and ethanol vapor. To avoid a naïve odor bias, reciprocal odor controls were run simultaneously. 30 min or 24 hours after training, flies were given a choice between unpaired and paired odors in a Y-maze (79) (Fig 6A). A preference index was calculated by subtracting the number of flies entering the unpaired odor vial from the paired odor vial, and dividing this number by the total number of flies tested (78). A conditioned preference index was calculated by averaging the preference index of the two reciprocal training sessions. 60 flies were used for each N=1, 30 flies for each reciprocal conditioning. N=20-26 for each strain per condition.

### Alcohol Metabolism

To investigate how the different *for* strains absorb and metabolize alcohol, we exposed flies to 60% vaporized ethanol (90:60 ethanol:air) for 10 min and measured internal ethanol concentration immediately after or 30 min after ethanol exposure. The internal ethanol concentrations were determined from whole fly homogenates of 50 flies per sample (78). To measure the ethanol concentration, flies were first flash frozen in liquid nitrogen, and then homogenized in 500 μl Tris-HCL (50mM, pH 7.5, Sigma), followed by centrifugation at 4°C at 14000 r.p.m. for 20 min. Next, the ethanol concentration of the supernatant was measured using an alcohol dehydrogenase-based spectrophotometric assay (Ethanol Assay Kit 229-29, Diagnostic Chemicals) (Fig 7A). To set this data in relation and thereby calculate the fly internal ethanol level, the flies volume was set to 2 μl (80).

### Statistical Analysis

For statistical analysis the locomotion activity data were split in four phases, baseline (0-300s), early ethanol (300-600s), late ethanol (600-900s), and recovery(900-1200s). All early ethanol, late ethanol and recovery data was normalized to baseline. Data for activity, velocity, touch and stop for each phase was averaged and analysed. Due to low sample size (n=8) in behavior tracking experiments, we used more conservative non-parametric tests for the statistical analysis. This increased rigor, ensured consistency in analyses, and allowed for easier comparisons between experiments. For comparisons between the three nonpaired strains, a Kruskal Wallis test followed by Dunnett’s multiple comparisons was performed. In order to analyze changes in behavior of each strain over time (paired data), a Friedmann test followed by paired Wilcoxon multiple comparisons was performed (n.s.=p>0.05, *=p<0.05, **=p<0.01, ***=p>0.001, ****=p<0.0001). The statistical analysis was performed in PRISM 7 (GraphPad).

## Results

### foraging does not affect open field behaviour in the absence of a stimulus

We first tested whether rovers (*for^R^*), sitters (*for^s^*) and sitter mutants with a rover background (*for^s2^*) differ in spontaneous behavior in an open field arena without an ethanol stimulus, which we termed ‘baseline’ (Fig 1A,B). We found all *for* strains showed the same levels of group activity (H(3, 24)=0.29, P=0.8)(Fig 1E,F), velocity (H(3,24)=4.63, P=0.1 (Fig 1G), touching (H(3,24)=0.08, P=0.9) (Fig 1I) and stopping (H(3,24)=0.4, P=0.8)(Fig 1J).

The pathlength of *for^s2^* flies was significantly shorter compared to *for^s^* and *for^R^* (Fig 1H) (H(_3,24_)=7.88, P=0.01). In general, flies showed a high percentage of stopping behavior (70%, Fig 1J), and only around 4% of the flies in each arena interacted with each other (Fig 1I).

### foraging affects locomotion in response to an ethanol odor

We then looked at whether *for* elicited foraging behavior in response to a non-intoxicating ethanol dose similar to that found in fermenting fruit (Levey, 2004). When the flies were exposed to a non-intoxicating 10 min exposure of 10:105 ethanol:air ratio, the percent of flies’ activity increased to up to 40% in *for^R^* flies (p=0.008) and only 25% in *for^s^* (p=0.008) and *for^s2^* (p=0.008) flies compared to baseline activity levels (Fig 2A). *for^R^* flies demonstrated an increased group activity throughout the 10-minute ethanol exposure whereas *for^s^* and *for^s2^* flies steadily decreased activity over time, resulting in a significantly lower activity at the end of ethanol exposure compared to *for^R^* flies (H(_3,24_)=7.96, P=0.02, *for^R^* vs *for^s^* (p=0.01), *for^R^* vs *for^s2^* (p=0.02)) (Fig 2B). Rovers travelled significantly longer distances during low dose ethanol exposure compared to sitters (0-5 min: H(3, 24)=13.92, P <0.0001, *for^R^* vs *for^s2^* (p=0.0006), 5-10min: H(3,24)=13.72, P=0.0011, *for^R^* vs *for^s^* (p=0.02), *for^R^* vs *for^s2^* (p=0.001)) (Fig 2C). *for^R^* flies also show a significantly higher velocity over the time of low dose ethanol exposure compared to *for^s^* and *for^s2^* flies (H(3,24)=18, P=0.0001, *for^R^* vs *for^s^* (p=0.002),*for^R^* vs *for^s2^* (p<0.0001)) (Fig 2D). Our values are comparable to a previous study that reported in the presence of food rovers walked 36.17cm/30sec (361.7cm/5min), and sitters walked 17.38cm/30sec (173.8cm/5min), suggesting that the low dose ethanol here represents a food-like odor to the fly (10, 81). Ethanol odor increased touching (*for^R^* (p=0.008), *for^s^* (p=0.02), *for^s2^* (p=0.04)) and decreased stopping (*for^R^* (p=0.008), *for^s^* (p=0.008), *for^s2^* (p=0.008)) in all strains (Fig 2 E,F). *for* did not significantly affect touching (H(_3,24_)=1.665, P=0.43) (Fig 2E). However, rovers stopped significantly more than sitters after five minutes of ethanol exposure (H(3, 24)=9.38, P=0.009, *for^R^* vs *for^s^* (p=0.03), *for^R^* vs *for^s^* (p=0.02)) (Fig 2F).

**Fig. 2.**
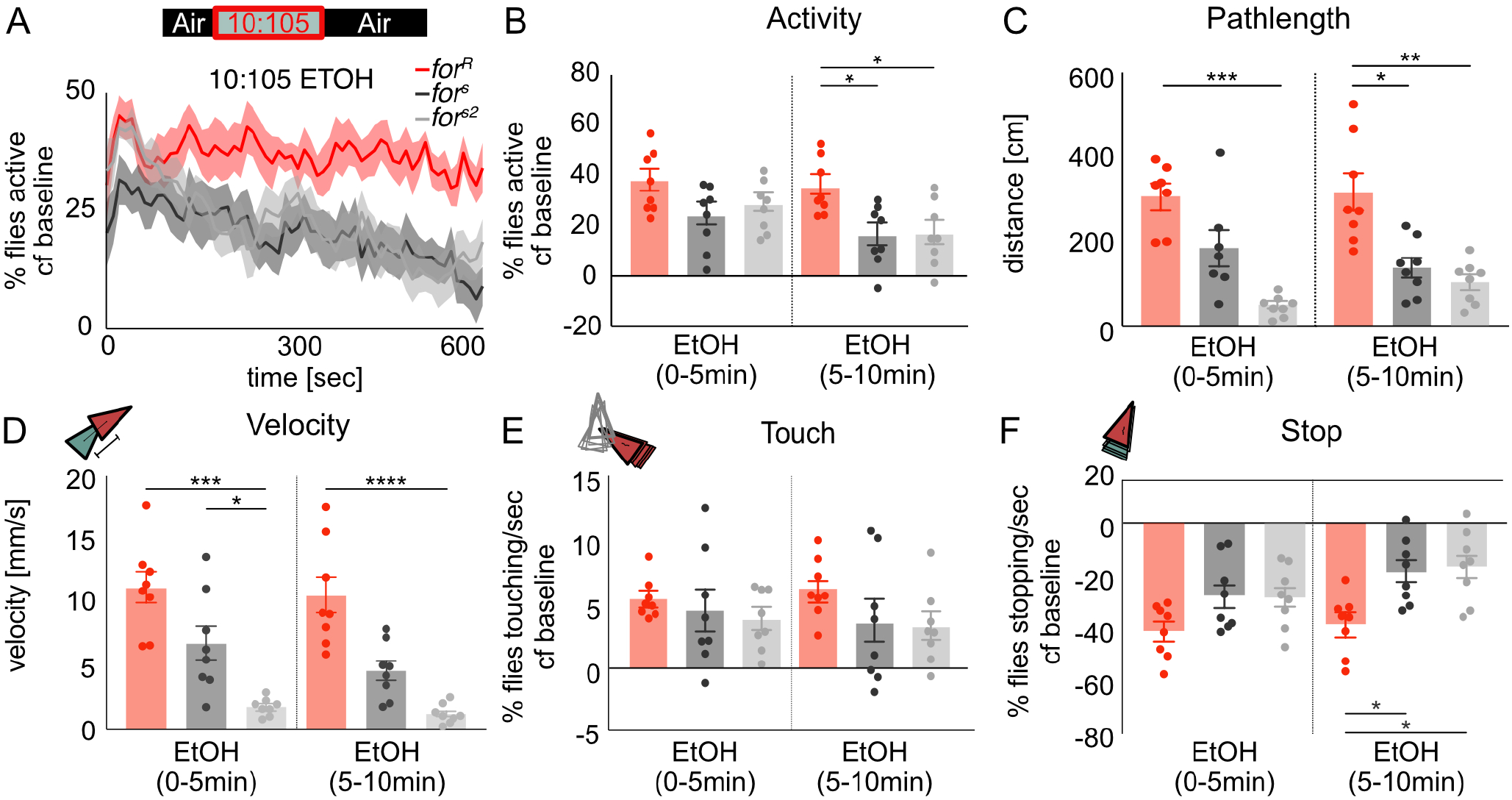
*for* affects activity in response to ethanol odor. (A,B) During the period of low-dose ethanol exposure, the activity for *for^s^* and *for^s2^* decreased, whereas the activity level of *for^R^* stayed the same. Lines depict mean+/- standard error. (B-F) Graphs show mean+/- standard error. (C) *for^R^* traveled significantly longer distances during low dose ethanol exposure compared to *for^s^* and *for^s2^* flies. (D) *for^R^* show a significantly higher velocity compared to *for^s^* and *for^s2^* flies. (E,F) Touching behavior increases for all strains compared to open field behavior whereas stopping behavior significant decreased to less than 60% of the flies stopping per second. *for^R^* stopped significantly less than *for^s^* and *for^s2^* flies. n.s.=p>0.05, *=p<0.05, **=p<0.01, ***=p>0.001, ****=p<0.0001)

### foraging does not affect behaviour post-ethanol odor stimulus

We hypothesized that once the ethanol odor stimulus was removed, all strains would recover to baseline behavior, resulting in no differences in behavioral measures between the three *for* strains. As predicted, there were no significant differences in group activity between rovers and sitters (Fig 3A) (*for^R^* vs *for^s^* (p=0.06), *for^R^* vs *for^s2^* (p>0.99)). Although there was a reduction in pathlength (H(_3,24_)=15.7, P=0.0004), and velocity (H(_3,24_)=15.61, P=0.0004) in *for^s2^* flies, and in stopping in *for^s^* flies (H(3, 24)=9.97, P=0.007), there were no consistent significant differences between *for^R^* and the two sitter strains in any of the metrics reported, so we do not attribute these differences to variation in *for* (Fig 3 A-F). There were no significant differences in touching between the three strains (H(_3,24_)=3.55, P=0.17) (Fig 3E).

**Fig. 3.**
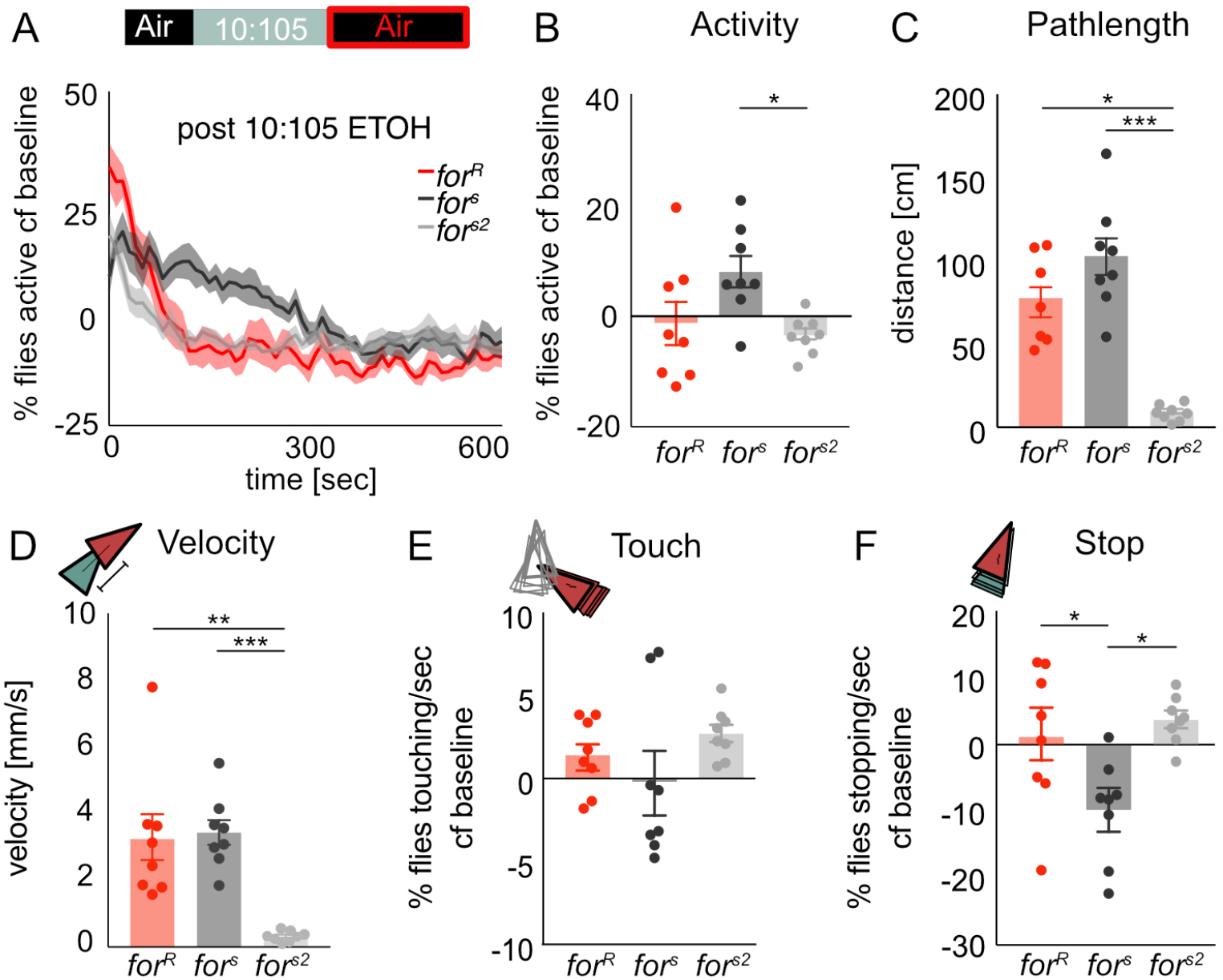
*for* does not affect recovery from ethanol odor exposure. (A,B) During presentation of humidified air following exposure to low-dose ethanol, the group activity returns to baseline activity with 300sec, with no significant differences between strains. Lines depict mean+/- standard error. (B-F) Graphs show mean+/- standard error.(C) The total distance travelled (pathlength) after low-dose ethanol exposure returns to baseline behavior in *for^R^* and *for^s2^* flies, whereas *for^s^* flies still show an increase pathlength compared to baseline. (D) The velocity after low-dose ethanol exposure returns to baseline behavior in *for^R^* and *for^s2^* flies, whereas *for^s^* flies still show an increase velocity compared to baseline., (E,F) Touch and Stop behavior recover within minutes of humidified air post ethanol odor. *for^s^* flies demonstrate less stopping than both *for^R^* and *for^s^* flies. Since there were no consistent differences between rovers and the two sitter strains in any of the metrics reported, we do not attribute these behavioral differences to variation in *for*. n.s.=p>0.05, *=p<0.05, **=p<0.01, ***=p>0.001, ****=p<0.0001

### foraging affects ethanol-induced increases in locomotion and touching

We next investigated how variants of *for* affected these behavioral metrics in response to a dose of ethanol that typically induces sustained increases in locomotor activity (Scaplen et al., 2019), by exposing the flies to a 50:65 ethanol:air mixture (43% ethanol vapor) for 10 minutes. Sitter strains increased group activity by 40% (*for^s^* (p=0.008), *for^s2^* (p=0.008) whereas rovers only showed 30% activity increase (*for^R^* (p=0.02) compared to baseline (Fig 4A). During ethanol exposure, *for^R^* flies showed a significantly lower activity than sitters (H(_3,19_)=10.9, P=0.004, *for^R^* vs *for^s^* (p=0.006), *for^R^* vs *for^s2^* (p=0.04)) (Fig 4B). However, *for^R^* did not show significantly less distance moved (p>0.9) or velocity (p>0.9) compared to *for^s^. for^s2^* flies showed significantly reduced pathlength (H(_3_, i9)=12.33, P=0.0002) and velocity compared to *for^s^* and *for^R^* flies (H(_3,19_)=9.47, P=0.004), but since *for^R^* flies did not differ from both sitter strains, we do not attribute these differences to variation in *for*. This dose of ethanol increased touching (*for^R^* (p=0.02), *for^s^* (p=0.03), *for^s2^* (p=0.04)) (Fig 4E) and decreased stopping (*for^R^* (p=0.03), *for^s^* (p=0.03), *for^s^*^2^ (p=0.04)) in all strains (Fig 4F). During early ethanol exposure, *for^R^* flies showed significantly more touching (H(_3,19_)=12.19, p=0.0002) than *for^s2^* (p=0.002), whereas this effect trended towards significance with *for^s^* (p=0.06). In contrast, *for^R^* flies showed significantly more touching (H(_3,19_))=6.53, p=0.03) than *for^s^* (p=0.04) but not *for^s2^* (p=0.2) during late ethanol exposure (Fig 4E). *for^R^* flies showed significantly more stopping (H(_3,19_)=9.17, P=0.005) than *for^s^* (p=0.01, p=0.04) but not *for^s2^* (p=0.1, p=0.9) flies during early and late high-dose ethanol exposure (Fig 4E)..

**Fig. 4.**
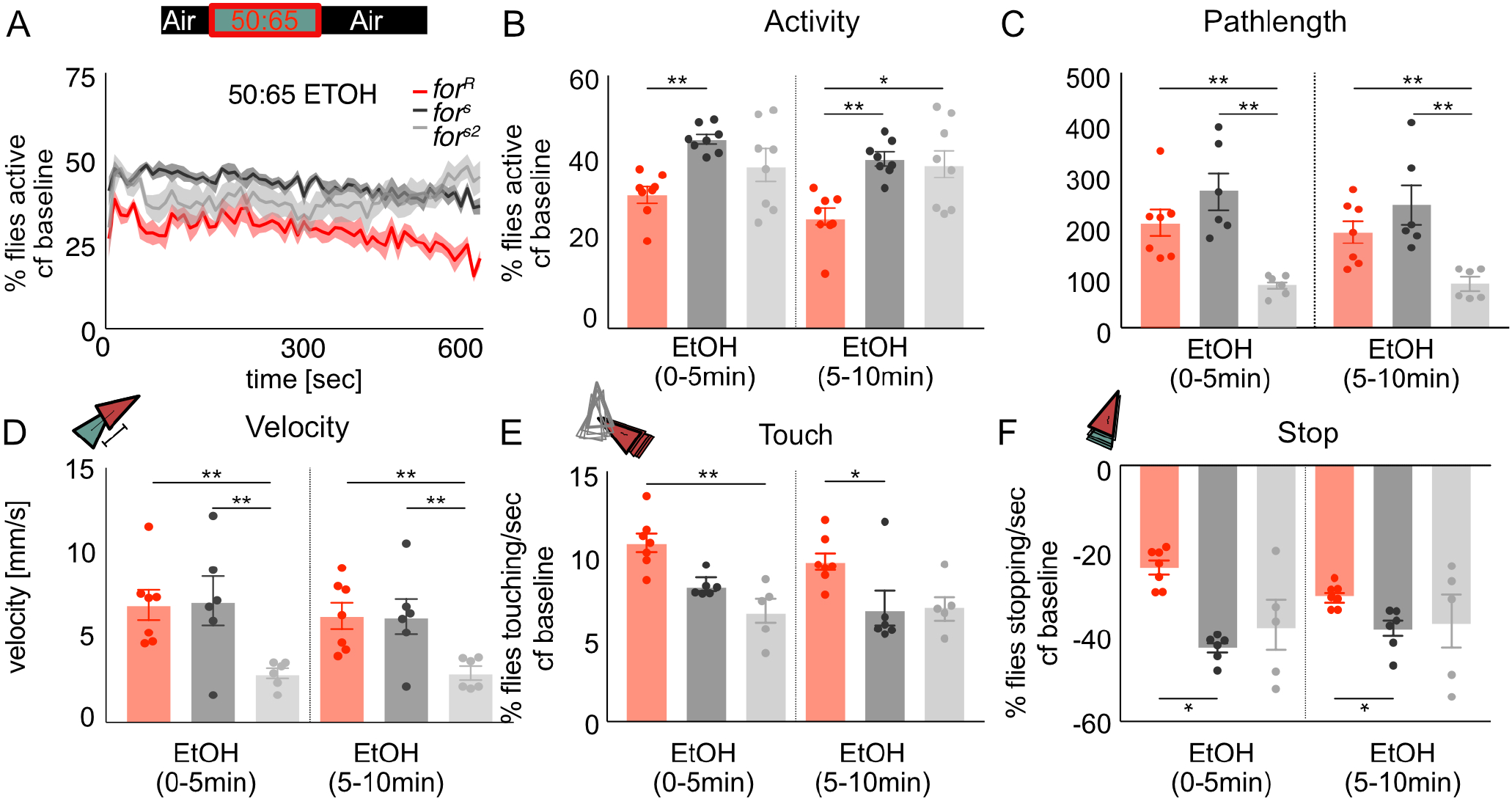
for affects behavioral sensitivity to the pharmacological properties of ethanol. (A) The percent activity in response to a high dose of ethanol increases to 35% for rovers and 45% for sitters. Lines depict mean+/- standard error. (B-F)Graphs show mean+/- standard error. (B) During high dose ethanol exposure the group activity of *for^R^* is significantly lower than *for^2^* and *for^s2^* flies. (C) *for^s2^* flies move a significantly smaller distance than *for^R^* and *for^2^* flies. (D) *for^s2^* flies are significantly slower than *for^R^* and *for^2^* flies. (E,F) All strains showed increased touching and decreased stopping during high-dose ethanol exposure. *for^R^* showed significantly more touching than *for^2^* and *for^s2^* (E) and significantly more stopping than *for^2^* and *for^s2^* flies in the first five minutes of ethanol exposure (F). n.s.=p>0.05, *=p<0.05, **=p<0.01, ***=p>0.001, ****=p<0.0001

### foraging affects recovery from the pharmacological properties of ethanol

We hypothesized that since *for* affected the percent of flies active during ethanol intoxication, it may also affect recovery from the pharmacological properties of ethanol. Indeed, significantly fewer rover flies were active compared to sitters after the offset of ethanol (H(_3,19_)=14.22, P=0.0008, *for^R^* vs *for^s^* (p=0.0009), *for^R^* vs *for^s2^* (p=0.02)) (Fig 5A,B). However, *for^R^* flies moved significantly more distance than *for^s2^* flies (p=0.04) but not *for^s^* flies (p>0.9) (Fig 5C). *for^R^* pathlengths returned to baseline within 5min of recovery (p=0.4), which was not the case for *for^s^* (p=0.03) or *for^s2^* (p=0.03). Similarly, *for^R^* flies moved significantly faster than *for^s2^* flies (p=0.03) but not *for^s^* flies (p>0.9) (Fig 5D). These activity and locomotion metrics suggest that the 50:65 ethanol:air dose did not sedate the flies, and that rovers recover from the pharmacological properties of ethanol faster than sitters.

**Fig. 5.**
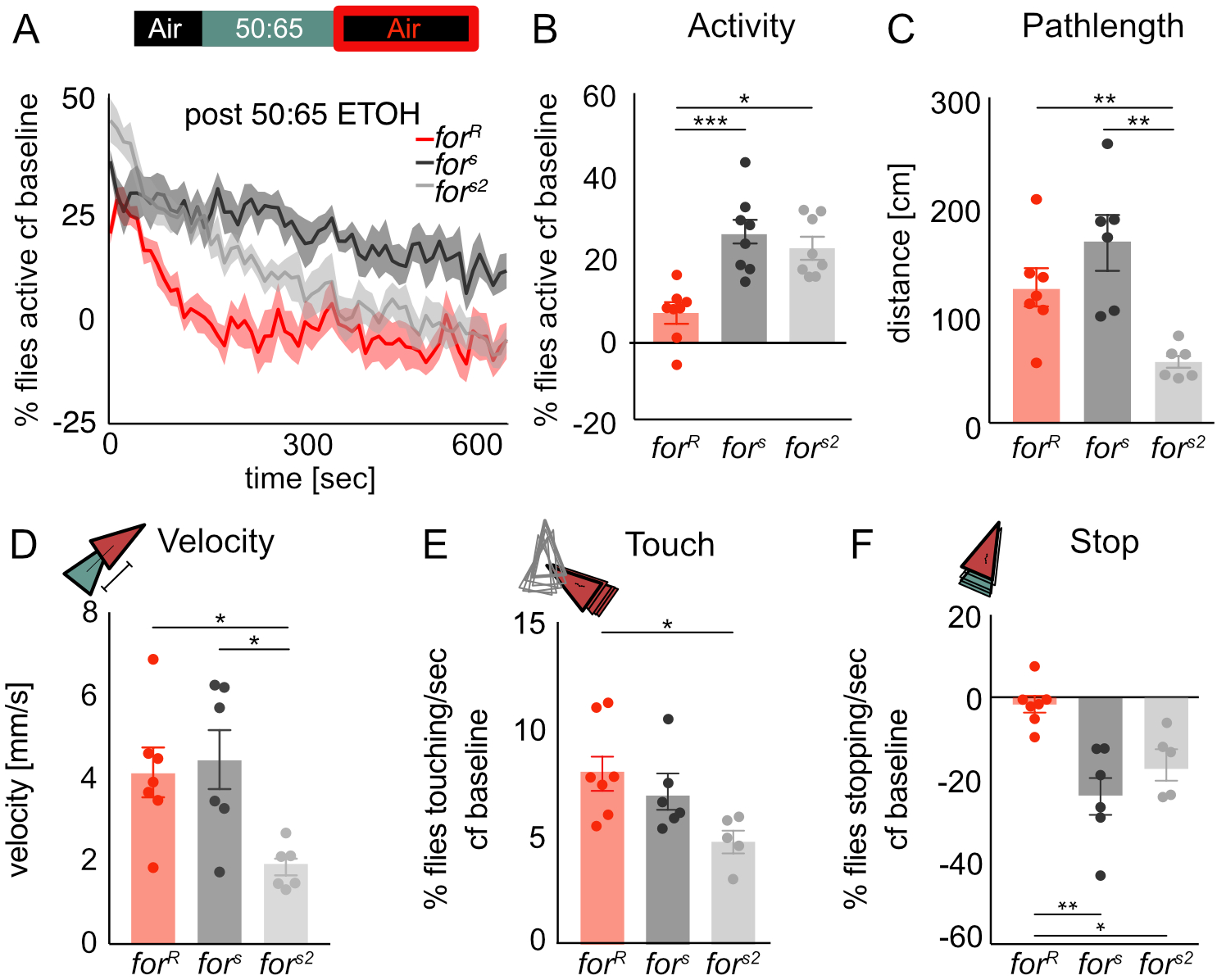
*for* affects recovery from the pharmacological properties of ethanol. (A) During exposure to humidified air following an intoxicating dose of ethanol, fly activity decreases more rapidly in *for^R^* than *for^s^* or *for^s2^* flies. Lines depict mean+/- standard error. (B-F) Graphs show mean+/- standard error. (B-F) Graphs show mean+/- standard deviation (B) Significantly fewer *for^R^* flies were active compared to *for^s^* and *for^s2^* flies. (C) There is significantly reduced (C) pathlength and (D) velocity in *for^s2^* flies compared to *for^R^* and *for^s^* flies (E) Touching behavior remained higher than base-line levels for all strains, and was statistically reduced in *for^s2^* compared to *for^R^* flies (F) *for^R^* stopping behavior returned to baseline levels whereas *for^s^* and *for^s2^* flies continued to show significantly less stopping compared to baseline. n.s.=p>0.05, *=p<0.05, **=p<0.01, ***=p>0.001, ****=p<0.0001

**Fig. 6.**
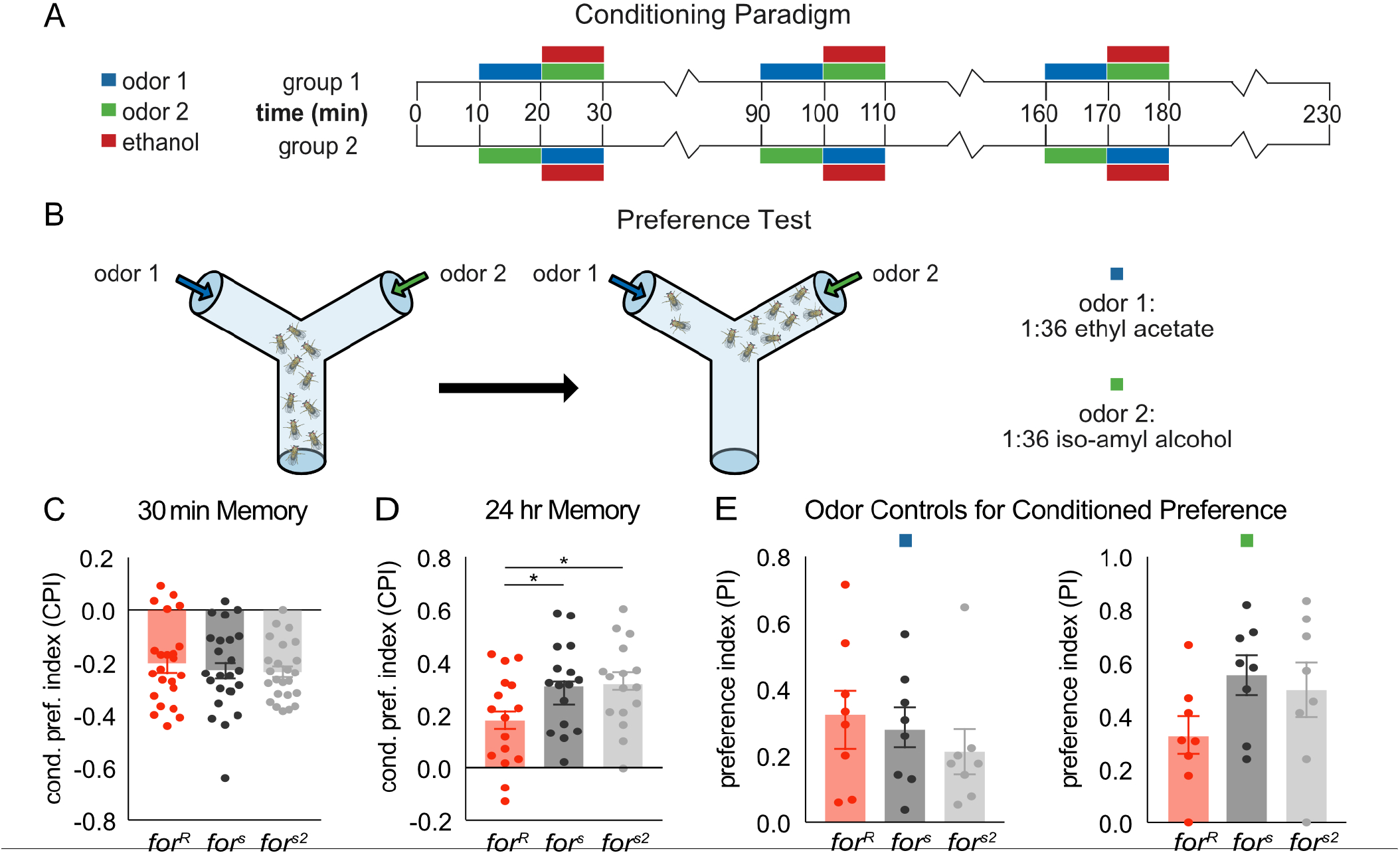
*for* affects memory for ethanol reward. (A) Flies received 3 training sessions with a 10 min exposure to one odor followed by a 10 min exposure to a second odor paired with 60% ethanol vapor. Training trials were spaced by 1 hour. A reciprocal group, in which the opposite odor was paired with ethanol, was run. (B) After the training flies were given the choice of the two odors. A preference index was calculated by subtracting the number of flies entering the Odor- vial from the Odor+ vial and dividing this number by the total number of flies. Conditioned preference index was calculated by averaging the preference indexes of the two reciprocal groups. (C) *for* does not affect conditioned aversion tested 30 min after training. (D) *for* affects conditioned preference tested 24 hrs. after training. *for^R^* flies show decreased memory for ethanol reward compared to for2 and *for^s2^* flies. (E) *for* does not significantly affect preference for the odors used in the conditioning procedure. n.s.=p>0.05, *=p<0.05, **=p<0.01, ***=p>0.001, ****=p<0.0001

**Fig. 7.**
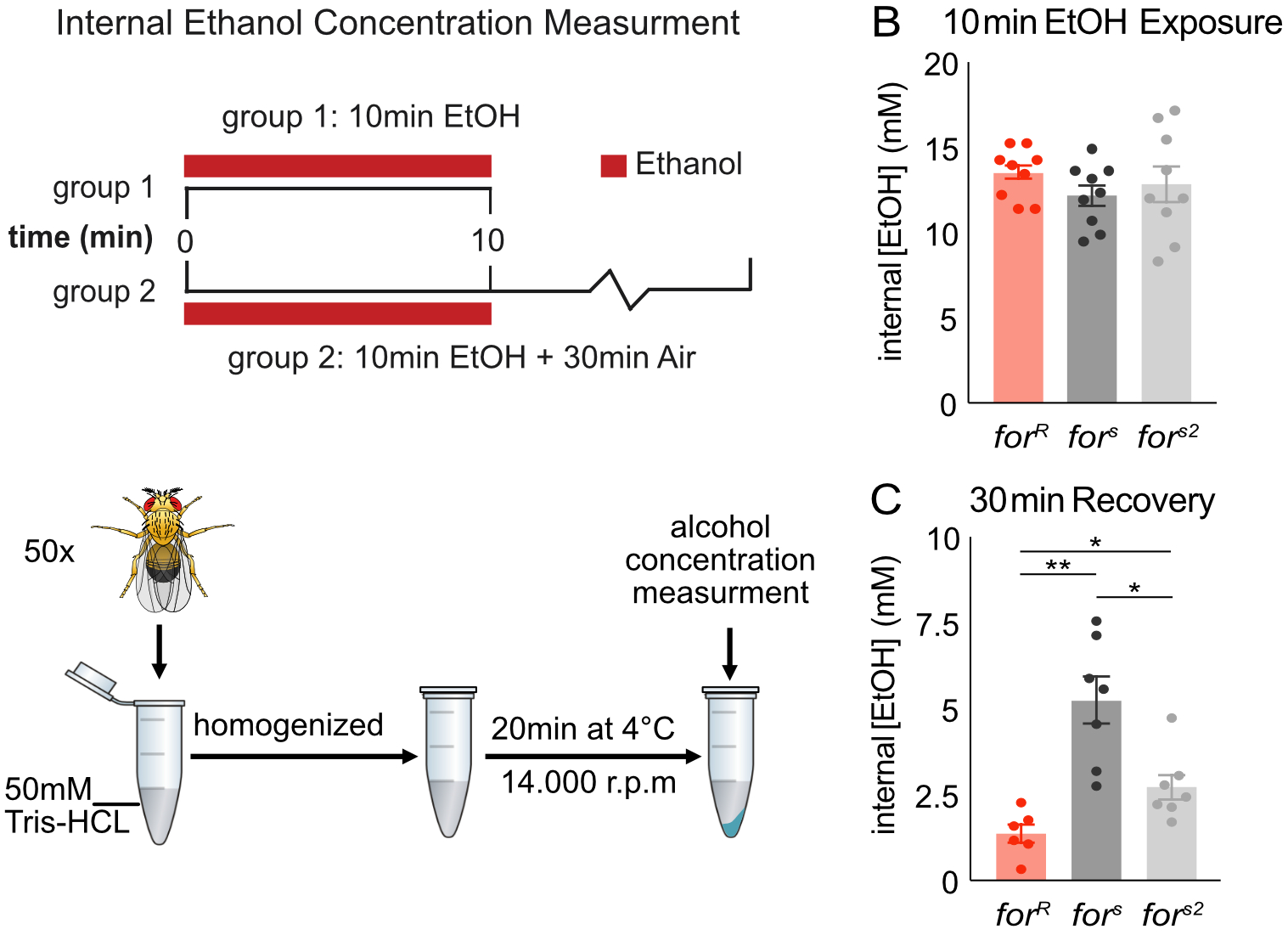
for affects ethanol metabolism. (A) Flies received a 10 min exposure to 60% ethanol in perforated vials in the training boxes used for ethanol memory. Flies from Group One were frozen immediately in liquid nitrogen, flies from group two are exposed to air for another 30min. Flies were homogenized in 500 μl of cold 50 mM Tris-HCl (pH 7.5, Sigma) and the homogenate was centrifuged at 14,000 r.p.m. for 20 min at 4 °C. Ethanol concentrations in supernatants were measured using an alcohol dehydrogenase–based spectrophotometric assay (Ethanol Assay Kit 229-29, Diagnostic Chemicals). To calculate fly internal ethanol concentration, the volume of one fly was estimated to be 2 μl. (B) *for* does not affect ethanol absorption. (C) *for* affects rate of recovery from ethanol exposure. *for^R^* flies metabolize ethanol faster than *for^2^* and *for^s2^* flies. n.s.=p>0.05, *=p<0.05, **=p<0.01, ***=p>0.001, ****=p<0.0001

Touching behavior remained higher than baseline levels for all strains (*for^R^* (p=0.02), *for^s^* (p=0.03), *for^s2^* (p=0.04)), and was significantly greater in *for^R^* flies than *for^s2^* (p=0.01) but not *for^s^* flies (p>0.9) (Fig 5E). *for^R^* flies stopping behavior returned to baseline levels (p=0.6) whereas *for^s^* (p=0.03) and *for^s2^* (p=0.03) flies continued to show significantly less stopping compared to baseline (Fig 5F). *for^R^* flies showed significantly more stops (H(_3,19_)=11.93, P=0.0002) than *for^s^* (p=0.003) and *for^s2^* (p=0.04) flies (Fig 5F).

### foraging affects reward memory for a cue associated with ethanol

We next hypothesized that the alcohol behavioral sensitivity differences between rovers and sitters would influence how *for* affects memory for an odor cue associated with ethanol (Fig 6A,B). Typically flies demonstrate aversive memory for ethanol when tested within 9 hours after training, and an appetitive memory when tested 15 hours or more after training (78). Since *for* has been shown to affect both short-term and long-term memory, we hypothesized that *for^R^* would show increased 30-minute aversive memory and reduced 24-hour appetitive memory compared to *for^s^* and *for^s2^*. No statistical differences were found between strains when flies were tested 30 minutes after training (H(3,_71_)=0.63, P=0.73) (Fig 1C). However, 24 hours after training, *for^R^* flies show reduced memory for ethanol compared to *for^s^* and *for^s2^* flies (H(_3,71_)=10.27, P=0.006, *for^R^* vs *for^s^* (p=0.03), *for^R^* vs *for^s2^* (p=0.002)) (Fig 1D). No statistical differences were identified in the ability of the strains to smell the odors used in the conditioning assay (IAA(H(3 24)=3.79, P=0.15), EA(H(3 24)=1.28, P=0.53)) (Fig 1E).

### foraging affects ethanol metabolism

Since *for* affects a number of metabolic phenotypes (4, 14, 82) we hypothesized that the observed behavioral differences between rovers and sitters may stem from the ability to metabolize ethanol differently. Therefore, we tested the internal ethanol levels right after ethanol exposure and after 30 minutes of recovery. We found that *for^R^* flies absorb ethanol at the similar rate (H(_3,27_)=1.87, P=0.39) (Fig 7B), but metabolize ethanol faster than *for^s^* and *for^s2^* flies (F(3,20)=13.6, P<0.0001, *for^R^* vs *for^s^* (p=0.003), *for^R^* vs *for^s2^* (p=0.02)) (Fig 7C).

## Discussion

We found that *for* affects alcohol induced locomotion (pathlength, velocity, activity, and stopping), social behavior (touching), cue-associated memory for the intoxicating properties of ethanol, and ethanol metabolism.

We first recapitulated previously published work demonstrating that rovers move more than sitters in response to a food stimulus. Rovers showed significantly longer path-length and moved faster compared to the sitter strains in response to a low-dose ethanol odor. This is consistent with previous work describing how rovers walk longer distances while searching for food whereas sitters tend to stay at one food source once they find it (12).

In contrast to these ethologically-relevant ethanol odor responses, we found that a pharmacologically-relevant ethanol concentration induces a sustained increase in the number of flies active within a group of sitters, but not rovers. However, rovers move similar distances and at the same speed as sitters under these conditions. They also touch more often, and stop less than sitters. This suggests that sitters are more sensitive than rovers to pharmacologically relevant stimulating doses of ethanol. This behavioral phenotype is consistent with the slower return to baseline levels after removal of an ethanol stimulus, and with a slower metabolism of ethanol in sitters.

Here we also showed that the foraging gene affects memory for the intoxicating properties of ethanol, since rovers show reduced preference for an ethanol associated odor cue 24 hours after the association. This is consistent with known roles of *for* in learning and memory. Rover larvae are better able to acquire and remember three but not eight odor-sucrose pairings compared to sitter larvae (82). Similarly, adult rover flies have better short-term olfactory memory, but worse long-term olfactory memory than sitters (28). This phenotypic difference is also seen in visual learning paradigms and is conserved in mammals (83, 84). Rovers also show higher retroactive interference, which occurs when the retrieval of previously learned information is less available owing to the acquisition of recently acquired information (27, 29, 30).

In the case of ethanol reward memory in *Drosophila*, we speculate that the difference in ethanol sensitivity between *for* strains alters perception of the intoxicating experience, causing reward to be processed differently in rovers and sitters. This may explain why *for* does not affect aversive short-term memory for ethanol, as this type of memory is not dependent on intoxicating concentrations of ethanol (79). Since *for* affects sensitivity to sucrose and other food substances, altering perception of the reward stimulus may be a more general mechanism through which PKG influences alcohol preference (3, 75).

Alternatively, as *for* also affects alcohol-induced increases in activity, this change in behavior could be affecting memory acquisition independent of reward perception. Notably, *for* would not affect the behavioral choice during the memory test since the flies are no longer intoxicated during odor choice (Fig 6A) and *for* does not appear to significantly affect preference for the odors used (Fig 6E).

Ethanol sensitivity has been associated with greater consumption and risk for developing an AUD (85–87). In human and rodents, sensitivity to the effects of alcohol intoxication is partially influenced by genes, whereas reduced sensitivity predicts the development of alcoholism (88–90). Thus, both heightened alcohol stimulation and reward sensitivity, and lower sensitivity to alcohol sedation robustly predicts more AUD symptoms over time in humans (86, 90). Studies in *Drosophila* recapitulate this, where genes influence the level of response to an intoxicating dose of ethanol (62, 91–94). Genome-wide association studies (GWAS) for alcohol sensitivity using the sequenced, inbred lines of the *Drosophila* genetic reference panel (DGRP) together with quantitative trait locus (QTL) mapping in an advanced intercross outbred population derived from sensitive and resistant DGRP lines, revealed 247 candidate genes affecting alcohol sensitivity, 58 of which, including the foraging gene, form a genetic interaction network (95).

Notably, our work did not demonstrate a consistent effect of for on touching. Although we observed a small trend where rovers demonstrated increased touching compared to sitters in the high-ethanol context (Fig 4E), this trend was not observed in any other behavioral contexts. Relatedly, for affected locomotion in the presence of a food odor in a similar way in our group assay as it did when flies are isolated in previously published studies (10, 12). Our lack of findings here was surprising as for affects social behaviors including aggregation behavior (96) and aggression (97), and influences behaviors that are dependent on a social context such as olfactory learning (98), and oviposition (99). Similarly, for orthologues influence social behaviors in other taxa including bees (100–102), wasps (103) and ants (104, 105). Our results failed to demonstrate an observed decrease in social dynamics in rovers compared to sitters that was recently observed in a similar assay (106). We speculate that sensory cues necessary for spontaneous social behaviors may have been obscured in our paradigm by a constant flow of hydrated air or odors through the behavioral chambers. This suggests that our behavioral paradigm was not optimal for identifying how for affected social behavior, or how social context could influence for-dependent behaviors.

Taken together, the work here contributes to a mechanistic understanding of alcohol sensitivity as indication for AUD by demonstrating how natural variation in metabolic phenotypes can impact behavioral response to an addictive substance. Our data predict that variants of *for* with lower PKG activity in other species will show increased ethanol sensitivity, and increased lasting ethanol preference.

This is consistent with results on the role of PKG in ethanol-induced behaviors from rodent models, suggesting the effects of PKG on alcohol behaviors are highly conserved. cGMP-dependent protein kinase type II (cGKII) knockout mice showed elevated alcohol preference in a 2-bottle free choice test (44), demonstrating that reduction in PKG is associated with increased alcohol preference in both mice and flies. Moreover, cGMP activates NO, which inhibits dopamine release in the striatum in rats resulting in decreasing reward response for alcohol (43, 49). These studies are consistent with rovers showing decreased ethanol preference, as they have higher PKG activity than sitters. Whether this is the mechanism through which variation in PRKG1 increases risk for alcohol abuse in humans remains to be seen (39, 40).

## ACKNOWLEDGEMENTS

We would like to thank Dr. Ulrike Heberlein (Janelia Research Campus) for helpful early discussions about the alcohol-associated preference and metabolism data. Thanks to Dr. Kristin Branson (Janelia Research Campus), Dr. Alice Robie (Janelia Research Campus), Dr. Mayank Kabra (Janelia Research Campus) and members of the Ctrax and JAABA google groups for ongoing assistance with Ctrax and JAABA. Thanks to members of the Kaun Lab and Dr. Kristin Scaplen (Bryant University) for providing helpful feedback on data visualization, analysis, and writing. Thanks to Dr. Marla Sokolowski for providing the for strains. This work was funded by the National Institutes of Health (R01AA024434 to K.R.K).

With gratitude to Dr. Marla Sokolowski:

Thanks especially to Dr. Marla Sokolowski (University of Toronto), whose guidance was integral to the conceptualization of this work. It is the people that we work with, including incredible mentors like Marla, that help shape us as scientists. Marla has been a huge inspiration to me (Karla), and more broadly an entire generation of behavioral geneticists. Her contributions have made an important and indelible mark on the field. I’m extremely grateful to her for her rigorous scientific training and continued patience, encouragement and immense knowledge. Marla is a lifelong mentor, and I consider myself incredibly fortunate to have the opportunity to conduct a study related to her research in my own lab. This work was completed with immense gratitude to Marla, for setting the stage and making it possible for generations of scientists to use foraging as an example of how natural genetic variation affects behavior. We all thank-you Marla, for being the amazing mentor and scientist that you are.

